# Prosculpt: Lowering the Barrier to Computational Protein Design

**DOI:** 10.64898/2026.06.25.732351

**Authors:** Federico A Olivieri, Alina Konstantinova, Neža Ribnikar, Nej Bizjak, Žan Žnidar, Kiyan Abel, Eva Rajh, Ajasja Ljubetič

## Abstract

Over the past decade, protein design has evolved from a specialized discipline into a broadly accessible approach for engineering and interrogating biological systems. Despite these advances, protein design continues to be a technically challenging task, often requiring knowledge of programming to be able to use and combine the different software packages. To address this challenge, we have developed Prosculpt, an easy-to-use protein design pipeline. Prosculpt integrates RFdiffusion for backbone generation, ProteinMPNN for sequence design and multiple structure-prediction platforms (AF2, AF3, Colabfold, Boltz2). Candidate designs are evaluated using customizable Rosetta-based scoring protocols. Each project is specified through a single configuration file, enabling users with minimal computational expertise to perform sophisticated protein design tasks without writing code, while also allowing advanced users to access the full capabilities of the underlying programs. Prosculpt supports a wide range of applications, including design of symmetric homo-oligomers, design of binders, motif scaffolding, partial diffusion and fixed-backbone sequence redesign. By combining these capabilities within a single, user-friendly platform, Prosculpt provides a practical entry point to modern protein design for both novice and expert users.

## Introduction

### Protein Design

In recent years, amazing progress has been made in the field of computational protein design^1^. Tools such as AlphaFold2^2^, ColabFold^3^, AlphaFold3^4^, ProteinMPNN^5^, RFdiffusion^6^ and others have found a wide range of applications in various fields, from de novo designed binders as therapeutics, to vaccine development and novel nanomaterials^7–11^. However, the complexity of tools used for each design step within a pipeline can present a challenge for users.

To address this, several automated workflows have recently been developed. These frameworks aim to streamline backbone generation, sequence design, and structure prediction into more accessible pipelines and lower technical barriers for end users with less computational experience. For example, ProtFlow^12^ and ProteinDJ^13^ provide structured environments for running design tasks with reduced manual scripting. Other tools such as BindCraft^14^ and BinderFlow^15^ focus specifically on protein–protein binder generation, integrating backbone generation and scoring into semi-automated systems.

These frameworks address the growing need to make protein design more accessible to a wider audience. However, the design steps supported by many of these tools remain restricted, often focusing primarily on binder design, and have limited application for other design problems such as motif scaffolding, design of symmetric assemblies or partial redesign of an existing protein.

Additionally, the degree of flexibility offered to users varies considerably, ranging from highly structured workflows (e.g. BindCraft^14^) with limited customization to more flexible systems that require scripting knowledge to fully exploit (e.g. PyRosetta^16^ or RosettaScripts^17^). Many workflows require users to install and configure multiple external dependencies. In addition, while steps such as backbone or sequence generation are becoming increasingly automated, scoring and evaluation of designs rely on metrics that are task-specific and require additional steps.

To address these challenges, we developed **Prosculpt**, a flexible pipeline that streamlines computational protein design. Prosculpt was designed to make advanced protein design methods accessible to researchers without specialized computational expertise while retaining the versatility of RFdiffusion-based workflows. It supports a broad range of design tasks, including binder design, motif scaffolding and symmetric oligomer generation (Fig. 1). Prosculpt integrates all stages of the design workflow, from backbone generation and sequence design to structure prediction and scoring, while allowing users to tailor each step through customizable settings.

**Figure 1:**
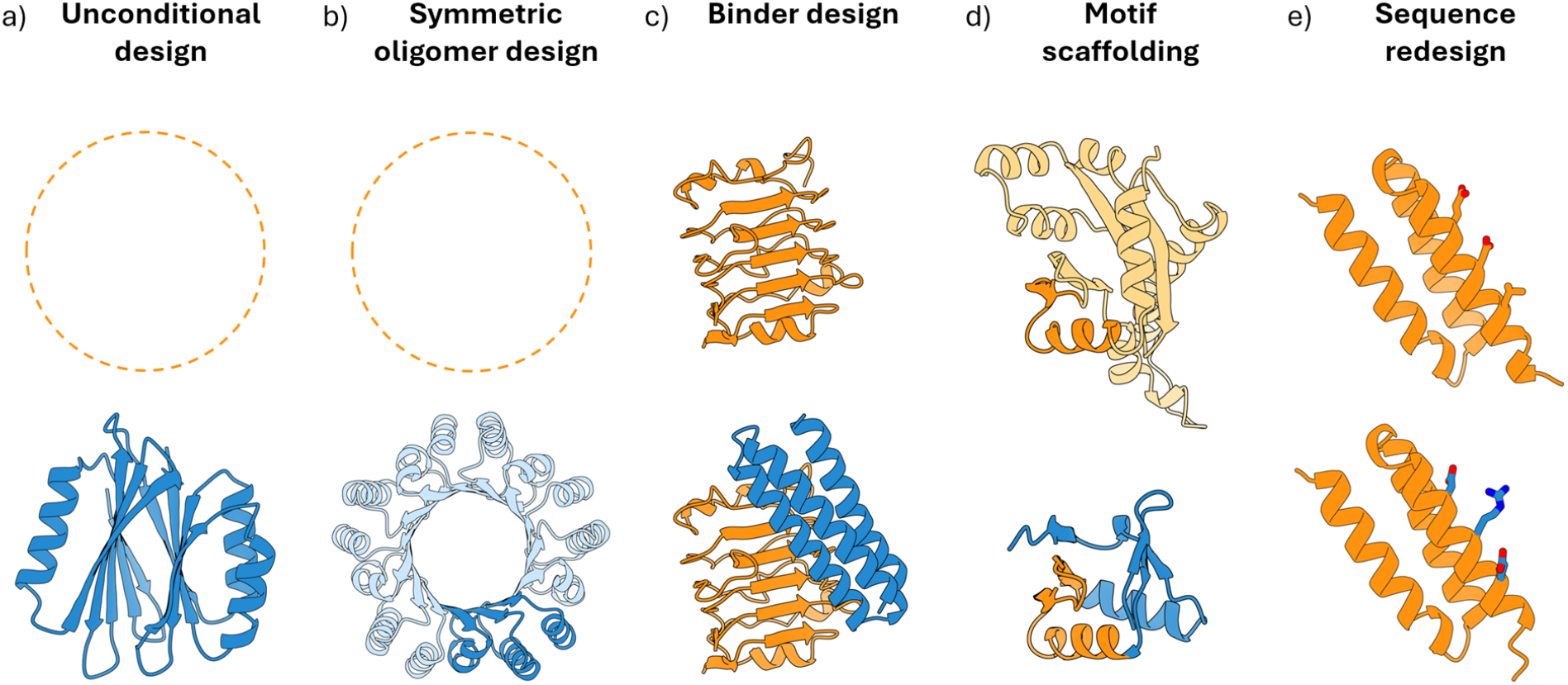
Representative protein design workflows supported by Prosculpt. Examples include **(a)** unconditional design, where a protein (blue) is generated without constraints (dashed orange), **(b)** symmetric oligomer design, where multiple identical units are designed at the same time, **(c)** binder design, where a protein (blue) is designed against a target (orange) **(d)** motif scaffolding, where a part of a protein (orange) is preserved an included in a new protein (blue) and **(e)** sequence-only redesign, where the backbone (orange) is fixed, but the sequence of the protein is redesigned (blue). These examples highlight selected capabilities and do not encompass the full range of supported applications.

Prosculpt additionally supports all RFdiffusion-based design modes, including partial diffusion and binder design against intrinsically disordered proteins, among others.

## Results

Prosculpt is a software package that automatizes the state of the art protein design pipeline consisting of backbone generation using RFdiffusion^6^, sequence design using ProteinMPNN^5^, structure prediction using one of several options and Rosetta for scoring (Fig. 2).

**Figure 2:**
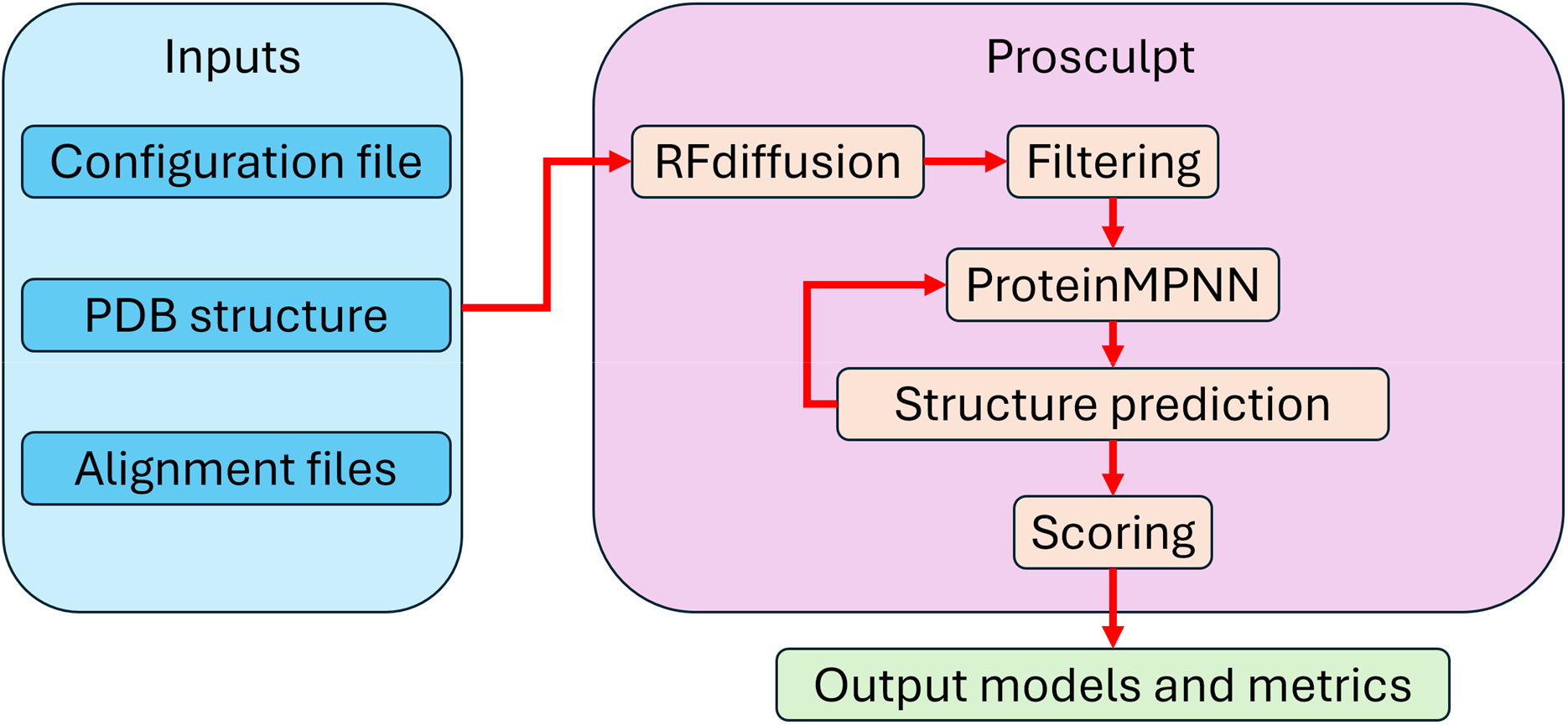
Overview of the Prosculpt design pipeline. User inputs (blue) consist of a configuration file and, where required, a protein structure (PDB file) and multiple sequence alignment information. Prosculpt then executes the complete design workflow automatically, including backbone generation with RFdiffusion, filtering, sequence design with ProteinMPNN, structure prediction and scoring (red arrows). The pipeline returns the final design models together with associated evaluation metrics (green), without requiring additional user intervention. A more detailed flowchart can be found in Supplementary Figure 1.

Prosculpt integrates these tools into a unified workflow by automatically converting the output of each program into formats compatible with subsequent stages of the pipeline. In doing so, it provides additional capabilities and customization options that are not available when the individual software packages are used independently. For example, in motif scaffolding it is possible to automatically only design the sequence of the newly generated backbone, while keeping the sequence of the rest of the protein fixed. It is also possible to generate metrics of only the newly generated parts of the protein, which is helpful in selecting the best designs.

Furthermore, Prosculpt supports seamless job recovery: if a run is interrupted it can be restarted, automatically resuming from the last completed stage and thereby mitigating the impact of common interruptions on shared computing clusters.

A Prosculpt run is configured through a single YAML-formatted file that specifies all parameters required across the pipeline. These parameters include, for example, the number of backbone structures to generate, the number of sequences to design per backbone, and the path to an input structure when required. Frequently used and required parameters are supported natively by Prosculpt through a streamlined interface. At the same time, users retain full access to the functionality of the underlying software, as arbitrary program-specific parameters can be passed directly to any stage of the pipeline. A full list of parameters can be found in Supplementary Table 1.

### Backbone design

The first step in the pipeline is backbone design using RFdiffusion. The required parameters for this step are the number of backbones to design and the contig string, which determines the length of sequence to generate and, if present, which residues to take from input structures. Additionally, the user may pass any settings to RFdiffusion, such as binding hotspots, number of diffusion steps or custom potentials. The Prosculpt repository contains examples of configuration files tailored to several different types of design jobs (e.g. Binder design, Partial diffusion, homooligomer design, etc) for users to adapt to their needs with minimal training.

After backbone generation, an optional filtering step can be applied, to conserve computational resources. Backbones that are not designable (i.e. contain large clashes or are only composed of one or two helices) can be eliminated at this step, saving the computational time need for sequence redesign and structure prediction. Filtering is implemented through Python scripts, termed *plugins*, each of which defines a filter_backbone function. This function accepts a PDB structure and user-defined arguments and returns a Boolean value indicating whether the structure satisfies the specified criteria. Prosculpt includes a recommended default plugin that filters designs based on the number of secondary-structure elements, removing proteins consisting of a single helix or a simple helix–loop–helix motif. Such architectures represent a common failure mode of RFdiffusion and are frequently associated with poor structural stability.

Users can easily develop custom plugins. These plugins are called in the configuration file, and multiple plugins can be called at once. Only designs passing all filters continue along the design pipeline. This approach separates filtering logic from the core pipeline implementation, allowing filtering criteria to change independently of the workflow itself and enabling users to adapt the pipeline to task-specific requirements without modifying the underlying codebase.

Prosculpt also allows users to bypass the RFdiffusion step entirely and perform sequence design directly on a user-provided backbone, either globally or over selected regions of the input structure. This mode is useful for fixed-backbone redesign, as all predictions and downstream metrics are generated and returned within Prosculpt.

### Sequence design

After all filters have been applied, ProteinMPNN is executed to generate sequences for each of the passing backbones. Residue positions eligible for redesign are obtained from the RFdiffusion TRB file. This is particularly valuable in motif scaffolding and binder design, where only newly introduced residues should be redesigned. Prosculpt also supports symmetric sequence design. This functionality can also be used to redesign existing symmetric assemblies (without generating a new backbone using RFdiffusion).

The required parameters in this step are the number of sequences to design per backbone, the backbone noise and the sampling temperature. The user can also choose whether to use the soluble or non-soluble ProteinMPNN model. Any other parameters accepted by ProteinMPNN can specified using the ‘pass_to_mpnn’ list of the configuration file.

### Structure prediction

After sequence design, AlphaFold2 (Colabfold), Boltz2 or AlphaFold3 can be used predict the structure of the new sequence. This is helpful for selecting the “best” designs, i.e. those that have the highest chance of working experimentally.

If no alignment information is provided, the designed sequences are used directly and modelled without multiple-sequence alignment (MSA) input. Alternatively, the user can provide an MSA file in a3m format for any chain in the original input structure. Using the information provided in these alignment files, Prosculpt generates partial alignment files, which retain all alignment information for natural protein fragments and add a “fake” MSA for the de novo designed part. The MSA consists of the *de novo* sequence copied along the rows.

Partial alignment files are particularly important when designing binders to natural targets or when scaffolding natural proteins. In the absence of a high-quality MSA, natural regions of the protein are poorly modelled in downstream prediction steps, which in turn reduces the reliability of subsequent filtering. In addition, this strategy reduces computational cost by avoiding large-scale alignment generation and repeated queries to external servers, thereby improving runtime efficiency.

The only required parameter for this step is the number of models to be created per sequence. In the case of AlphaFold2 (Colabfold), each structure is created by a specific trained model. Because some of these models are better for each specific tasks than others, the user can specify the models to use. For Boltz2 and Alphafold3, the number of structures represents the number of times a particular sequence is predicted to obtain a broader sampling.

When designing binders or symmetric assemblies, users may optionally generate models of individual binders or protomers in isolation, omitting the remaining chains, to evaluate intrinsic fold stability prior to complex formation.

Another option provided by Prosculpt is the ability to iterate using the predicted output structures as input for ProteinMPNN to refine the designs. Such iterative flexible backbone design improves sequence sampling on successful folds (Fig. 2)

### Scoring of designs

For each model created, Prosculpt compares the RFdiffusion model with the predictred structure. Standard metrics include RMSD and pLDDT, computed over multiple residue subsets: the full model, the “sculpted region” (residues that were de novo designed or modified from the input structure), “fixed chains” (chains that are unchanged from the reference structure) and “motif region” (non-redesigned residues within redesigned chains). These metrics are difficult to calculate without the integration between RFdiffusion and ProteinMPNN that Prosculpt provides.

If users choose to model binders in the absence of their target or operate in symmetry mode, the pLDDT and RMSD are calculated by comparing the prediction of a single chain with that of a single protomer extracted from the full model. Designs that maintain high pLDDT and low RMSD in the isolated state are typically better behaved and more likely to adopt the intended structure.

Prosculpt also calculates physics-based Rosetta metrics. The defaults are rg (radius of gyration), SAP (spatial aggregation propensity, representing the solvent-accessible hydrophobic surface area of a protein^18^) and charge of each design. These metrics are merged with the structural evaluation scores to generate the final scoring output. The scoring framework is designed to be modular, allowing users to modify the scoring script or implement custom metrics without altering the Prosculpt codebase. This provides flexibility for advanced users while maintaining a robust default configuration for non-specialist users.

Each design is assigned an ID, composed of 4 numbers separated by periods. These numbers correspond to task, backbone design, ProteinMPNN sequence and predicted structure respectively, allowing for unique identification and traceability of designs. After scoring is finished, all final models are pooled into one output folder and renamed to include their ID as well as metrics in their filenames to simplify manual selection and review.

### Getting started

Detailed installation instructions are available on the GitHub page. Prosculpt includes a collection of examples covering the most common design tasks in the **Examples** subfolder, including binder design, sequence redesign, partial diffusion, symmetric design, and others. Each example provides a configuration file together with the required structure and alignment files. Users can quickly get started by selecting the example that best matches their application and modifying the relevant parameters.

A Google Colab notebook for Prosculpt, featuring an intuitive graphical user interface and requiring no local installation, is also available (<u>Prosculpt_Colab.ipynb</u>). The interface is show in Supplementary Figure 2. Users can specify most Prosculpt parameters, including the input structure, the RFdiffusion contig, the number of RFdiffusion and ProteinMPNN designs, and the number of structure prediction models to generate. Sampling temperature, backbone noise, and symmetry constraints can also be defined, and advanced features such as partial diffusion and sequence inpainting are available. Users can perform a test run on the Google Colab platform. The resulting configuration file can be downloaded and directly transferred to a local Prosculpt installation for production runs.

### Application: design of C10 oligomers

A key advantage of Prosculpt compared to other protein workflow design libraries is the ability to design symmetric assemblies. Here, we demonstrate its utility in a challenging design problem by targeting C10 symmetric oligomers, that is proteins with 10 identical subunits arranged around a single axis of rotation. Notably, only a single de novo C10 oligomer has been reported to date^19^ underscoring the difficulty of the design problem.

We set out to generate symmetric oligomers with C10 symmetry. To increase backbone diversity, we systematically varied the protomer length and selected three design sets comprising 60, 90, and 120 residues. For each group, approximately 500 backbones were generated with RFdiffusion, then 10 sequences per backbone were generated with ProteinMPNN and the structures predicted with AlphaFold2 (Colabfold). Additional metrics used for design selection were calculated with Rosetta^20^, including charge, number of unsatisfied hydrogen bonds, SAP, Rosetta energy score (see Supplementary Table 2 for the full metric set). Designs were then ranked based on a combination of pLDDT and RMSD calculated between the AlphaFold2 model and original backbone generated with RFdiffusion.

Comparison of designs with different protomer sizes reveals a systematic decline in structural quality as complex size increases (Fig. 3). The *in silico* success rate (defined as the fraction of designs with pLDDT > 90) decreases from 7% for 60-residue protomers to 0.4% for 120-residue protomers, accompanied by an increase in backbone-to-AF2 model RMSD from 0.5 Å to 1.3 Å. These findings underscore a current limitation of RFdiffusion, whereby *in silico* design performance declines substantially for assemblies exceeding ∼400 amino acids in total length^6^.

**Figure 3:**
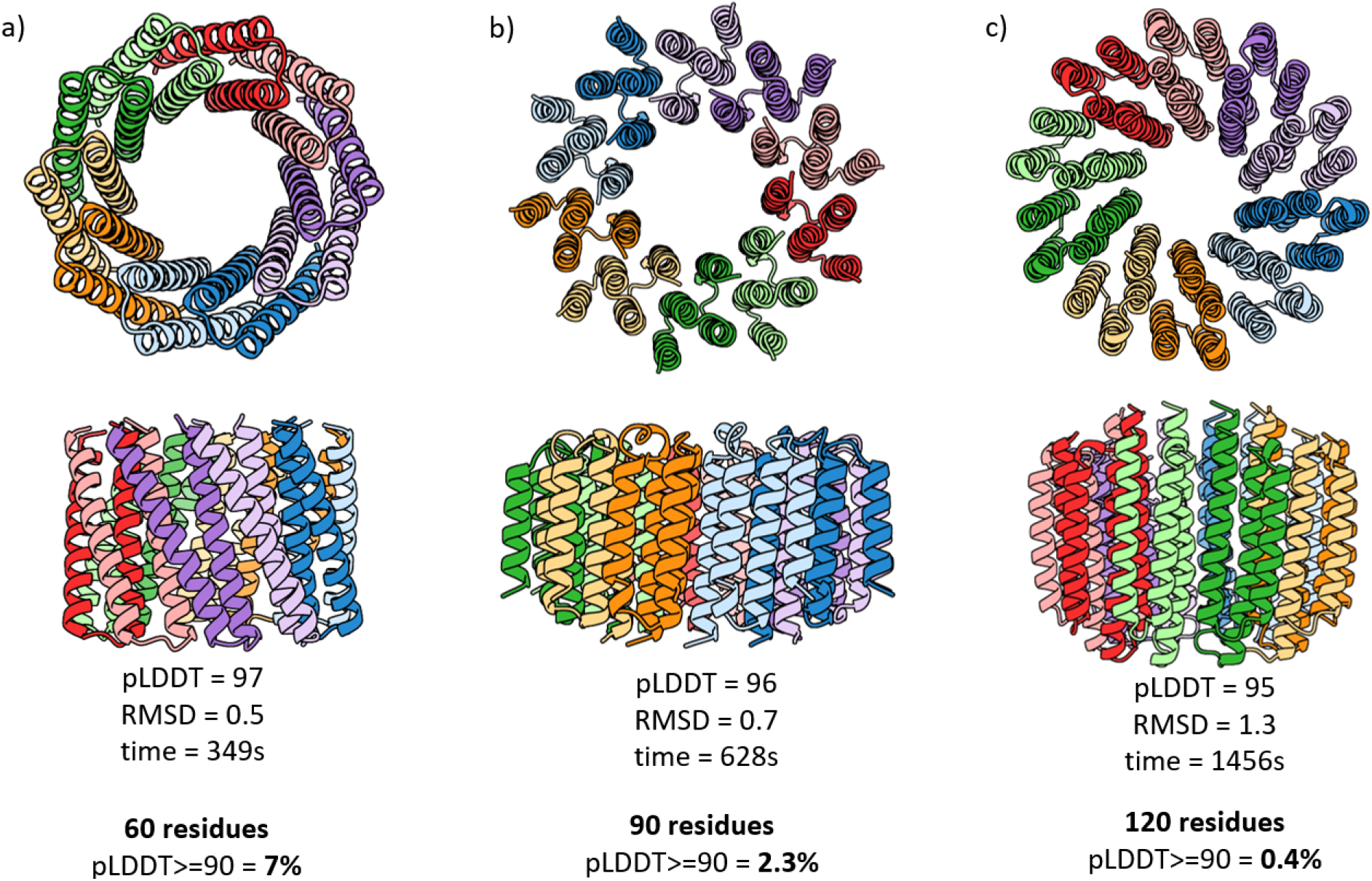
In silico success rates as a function of complex size. **(a)**C10 homo-oligomers, 60 residues per protomer. The representative design with the highest pLDDT is shown. Reported are the mean pLDDT, RMSD relative to the RFdiffusion backbone model, and total runtime for a single design. The *in silico* success rate (defined as fraction of designs with pLDDT ≥ 90) is indicated below. **(b)**Protomer designs of 90 residues, with analogous metrics as in (a). **(c)**Protomer designs of 120 residues, with analogous metrics as in (a). Across all panels, increased complex size is associated with reduced in silico success rate, increased structural deviation from the design model and longer run times.

A key objective of our design strategy was to generate a structurally diverse set of oligomeric assemblies for downstream experimental evaluation of alternative geometries. Among the highest-ranking designs, most monomers comprise 60 residues (Fig. 4), yet substantial variation is observed in both overall architecture and secondary-structure composition. The top backbones span fully α-helical forms (e.g., 7.7.2.1) to mixed α/β architectures with near-equal secondary-structure content (e.g., 4.43.8.1), while radii vary from 1.5 to 2.6 nm. Collectively, these results demonstrate that Prosculpt enables unconditional generation of symmetric protein assemblies with high in silico success rates and broad structural diversity.

**Figure 4:**
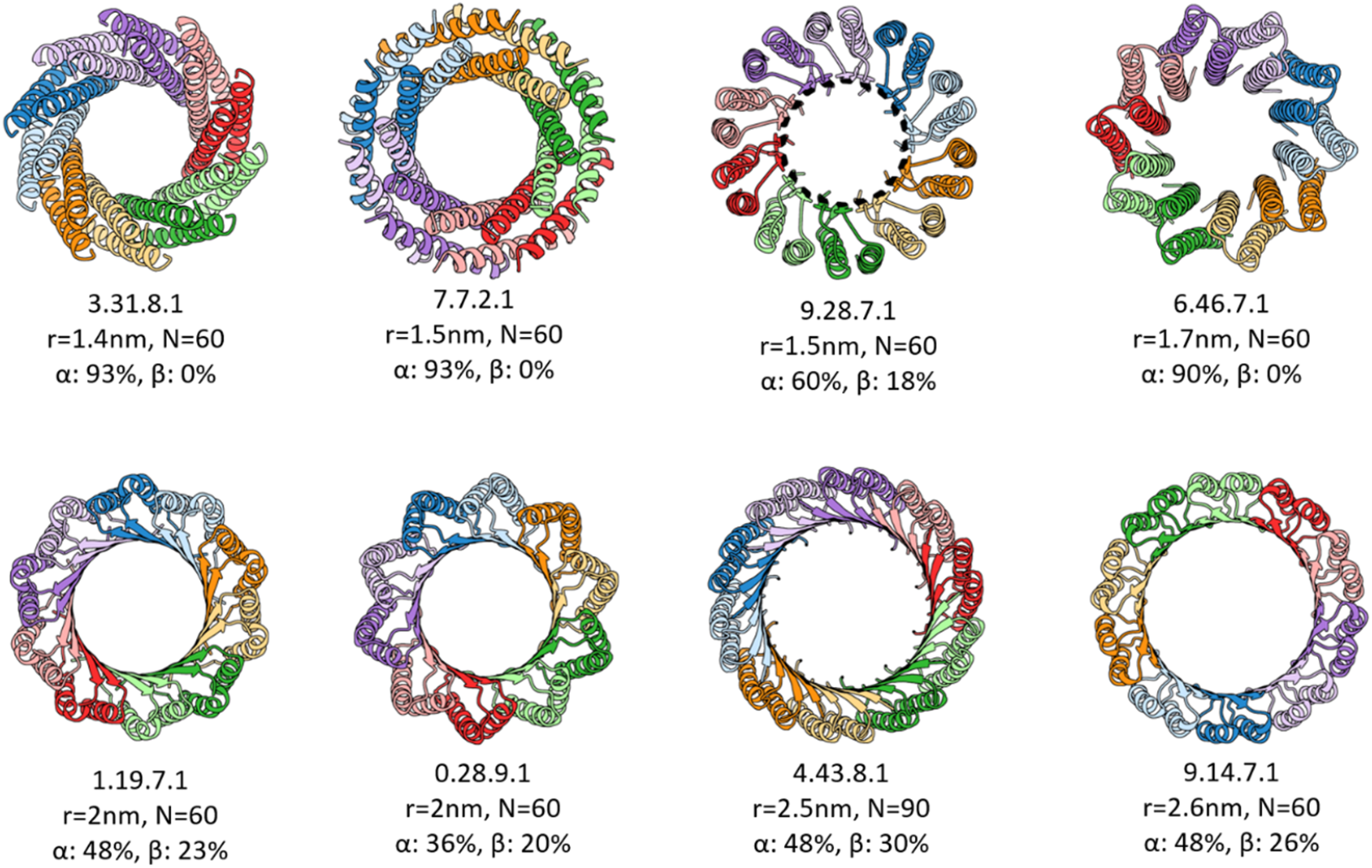
Backbone variability among top-ranked C10 oligomer designs. Each protomer is displayed in a distinct colour; four-letter design identifiers are indicated alongside structural annotations, including pore radius (r), number of amino acids per protomer (N), and the fractional α-helical and β-sheet content. The selected designs exhibit substantial variability in secondary-structure composition and pore geometry, spanning fully α-helical architectures to hybrids with nearly equal proportions of α-helices and β-sheets. Pore diameters range from 1.5 to 2.6 nm.

## Discussion

*De novo* protein design has the potential to become a routine tool in biochemistry laboratories for probing biological systems of interest. Prosculpt was developed to accelerate the adoption of de novo design by enabling users to initiate protein design workflows within minutes through a single configuration file. A suite of worked examples covering diverse applications further lowers the barrier to entry, allowing users to adapt pre-existing scripts to their own problems.

Multiple pipelines and workflows have been developed recently. BindCraft^14^ has a similar one-file configuration design as Prosculpt but can only perform binder design which it does through AlphaFold hallucination rather than diffusion. Ovo^21^ includes a graphical web interface, a custom quality control module called proteinQC which compares designs to reference values, but does not support many workflows and is also focused on binder design. ProteinDJ^13^ includes filtering steps not only after fold design but also after sequence design and model prediction, allowing for more granular control as well as an auxiliary tool to automate parameter screening.

Prosculpt, on the other hand, allows for any design task supported by RFdiffusion (Fig. 1). Notably, it is, to our knowledge, the only framework that natively supports design of symmetric oligomers, multi chain homo and heterodimers, partial diffusion and binders to targets with multiple chains. The platform further enables user-defined scripts for metric evaluation and backbone filtering, providing flexibility in defining scoring functions and selection criteria, which are often highly specific to the underlying design problem.

As protein design moves from a specialised computational discipline toward a routine experimental capability, workflow robustness and accessibility become increasingly important, particularly for laboratories without dedicated computational expertise. In our laboratory, Prosculpt has been applied to a range of design tasks, including binder engineering, symmetric homooligomer design, sequence redesign, terminal extensions, rigid fusion of protein components, and related applications. We anticipate that such workflow-oriented design tools will be broadly useful to researchers across the life sciences.

## Materials and Methods

### Prosculpt

Prosculpt was built in python 3.12. It is composed of two main files. The first is the Prosculpt library itself, prosculpt.py, which contains most functions regarding parsing of inputs and outputs of each step in the pipeline. The second is the prosculpt_run.py script, which serves as the entry point into the program. It uses hydra^22^ to parse the Prosculpt configuration files and the user-defined options and it calls each individual program in the pipeline accessing the Prosculpt functions in-between. Other important files are slurm_runner.py, which automatizes the parsing of slurm options from configuration files as well as the parallelisation over multiple slurm tasks, and the scoring script.

### Design of C10 homo oligomers

For the first round of designs, 14847 backbones were generated. Backbones were generated unconditionally, with monomer length set to 60, 90 and 120 residues, with 4999, 4998 and 4850 designs in each group, respectively. For these runs, guiding potentials were enabled, with the following parameters: type was set to olig_contacts, weight_intra to 1, weight_inter to 0, olig_intra_all and olig_inter_all set to True, and guide decay to quadratic. To increase backbone variability, we ran another batch of designs with monomer length of 60 residues and guiding potentials disabled.

For each backbone, 10 sequences were generated with ProteinMPNN, with sampling temperature set to 0.1 and backbone noise to 0. Each sequence was then validated with AlphaFold2. Metrics such as rg, charge, sap score were calculated with scoring_script.py as a part of Prosculpt pipeline, additional metrics (such as unsatisfied hydrogen bonds, Rosetta score, shape complementarity) were calculated with PyRosetta using an external python script (<u>extra_rosetta_metrics.py</u>).

Designs were initially filtered based on pLDDT and RMSD, with the cutoff pLDDT of 90, then the best designs were selected based on the combination of pLDDT, RMSD, number of unsatisfied hydrogen bonds, Rosetta energy score, as well as manual inspection of structures.

The comparison between *in silico* success rates for oligomers (Fig. 3) was made with guiding potentials enabled. Backbone variability is showcased using the best selected designs from both batches, with and without guiding potentials (Fig. 4). Pore radius for selected oligomers was measured using Pymol, calculated as half the mean Cα–Cα distance between equivalent pore-facing residues on opposing chains. Total runtime for a single design was calculated by dividing the runtime for the whole design batch by the total number of designs (500 designs per batch). The runtime was defined as a difference between the start time of first and the last steps of the design pipeline (RFdiffusion and scoring_script.py).

The configuration files used are available in <u>Examples/C10-oligomers</u> folder of the repository.

## Acknowledgements

This work was supported by the following funding sources: Slovenian Research and Innovation agency (ARIS) N1-0488, N1-0323, P4-0176 to A.L. The authors also thank the national computing cluster (VEGA) for GPU time through the project S25O02-12 awarded to A.L.

## Author Contributions

F.O. implemented partial alignments, developed the data transfer among modules, wrote the examples, implemented AF3 and Boltz2 and performed code maintenance. A.K. developed the symmetric oligomer design module of Prosculpt. N.B. wrote the initial version of Prosculpt, while Ž.Ž. contributed code development. K.A. developed the backbone filtering system. E.R. beta tested the software and contributed to its further development. A.K. and N.R. designed the oligomers.

A.L. conceived and directed the research, contributed to software development, and acquired funding. F.O., A.K., and A.L. drafted the initial manuscript. All authors reviewed, revised, and approved the final manuscript.

## Code and Data Availability

Prosculpt is distributed as an open-source software package available at https://github.com/ajasja/prosculpt/. Detailed installation and usage instructions are provided in the project documentation.

Models presented in Figure 3 and Figure 4 will be available at Zenodo at the time of publication.

## Supplementary Material

**Supplementary Figure 1:**
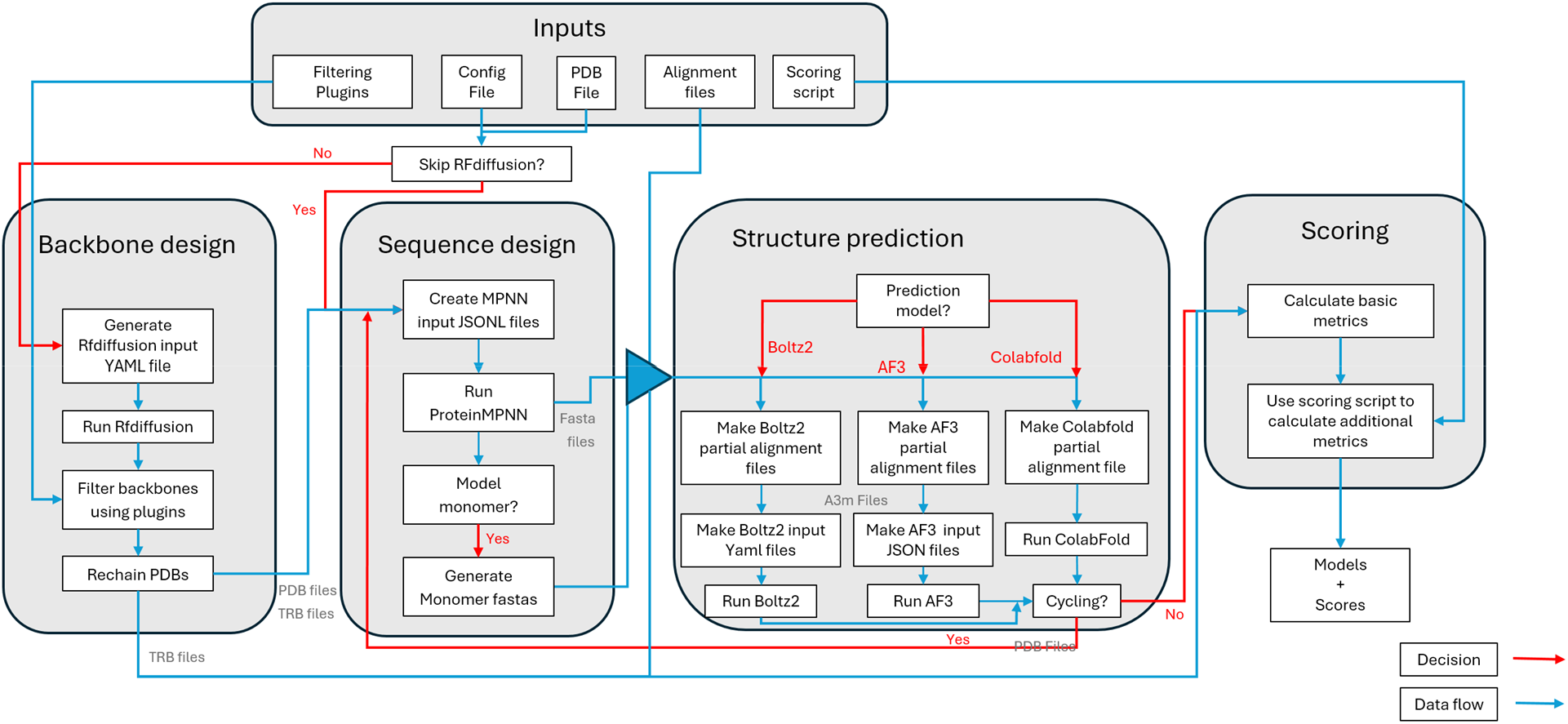
Flow diagram of Prosculpt. Blue lines denote data flow, while red lines denote alternate paths depending on configuration.

**Supplementary Figure 2:**
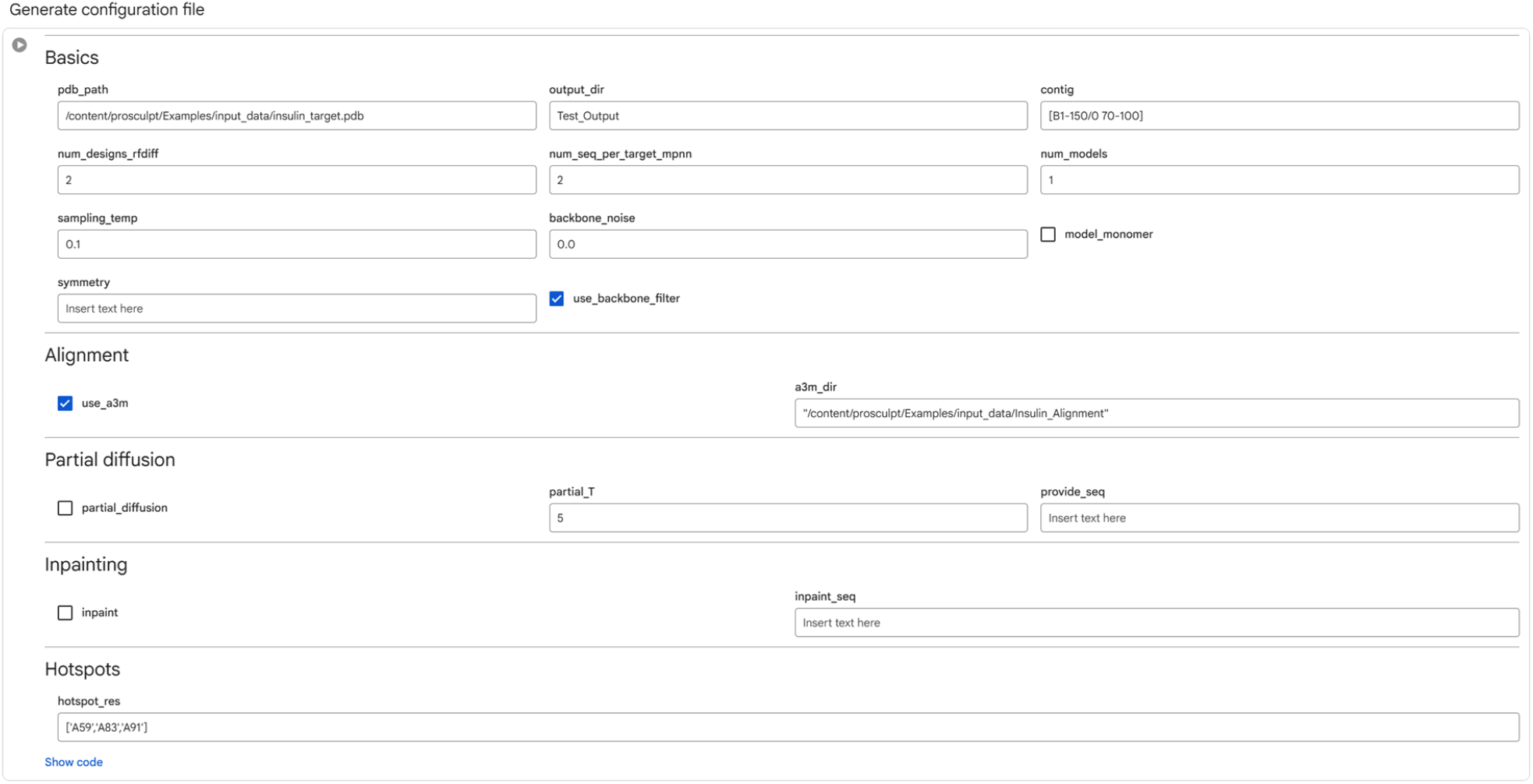
ProSculpt graphical user interface deployed in Google Colab. The interface provides an interactive, web-based environment for configuring and executing Prosculpt protein design workflows. Users can define input structures, the number of RFdiffusion and ProteinMPNN designs and the number of structure prediction models that are generated. Sampling temperature, backbone noise, and symmetry constraints can also be specified. Additional modules enable optional alignment using multiple alignment inputs, as well as advanced functionality for partial diffusion, sequence-conditioned design, and inpainting. Residue-level control can be specified through hotspot selection to guide design of binders. The configuration is automatically compiled into a single configuration file for downstream model execution. The configuration file can also be downloaded.

**Supplementary Table 1:**
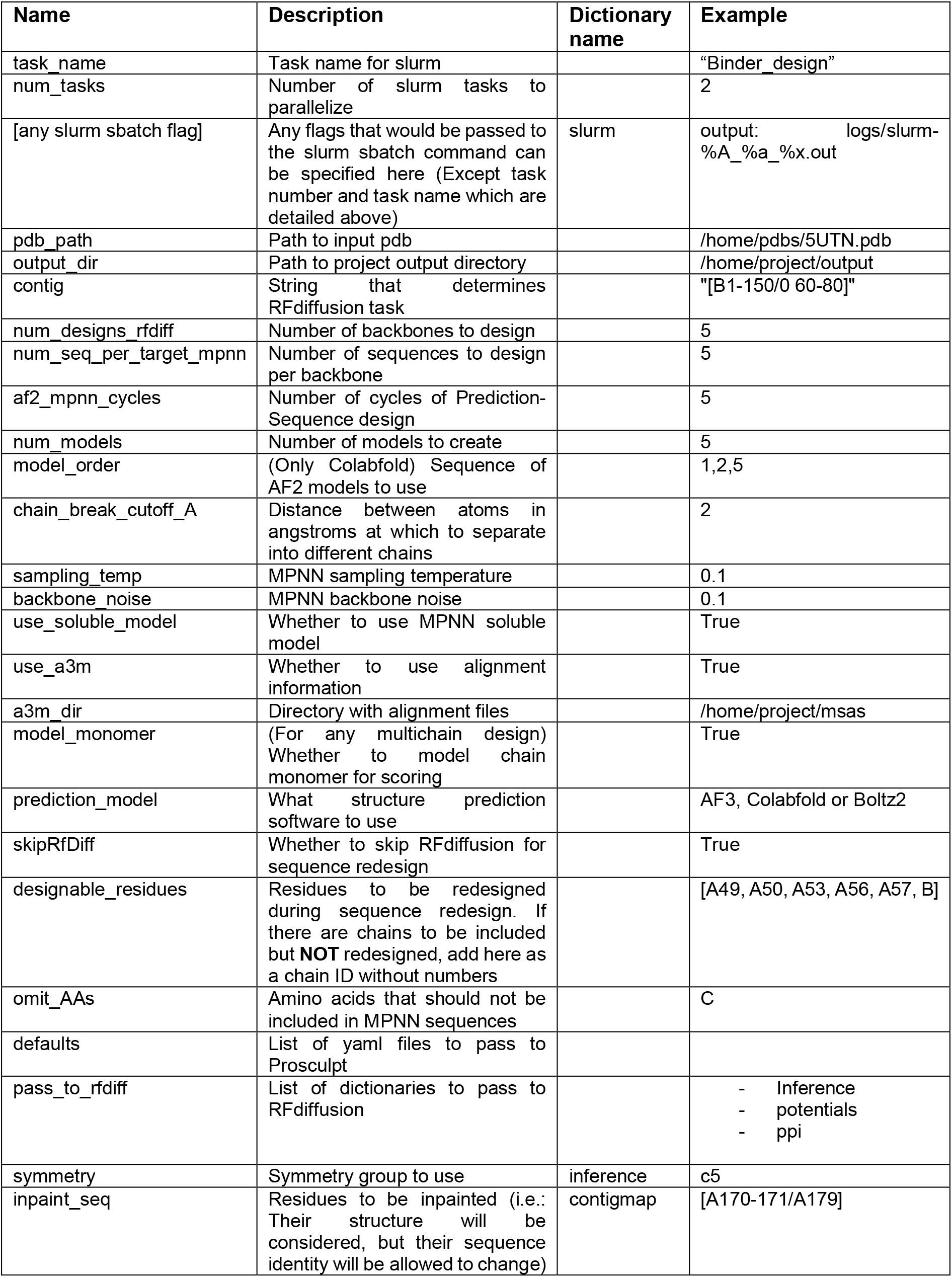
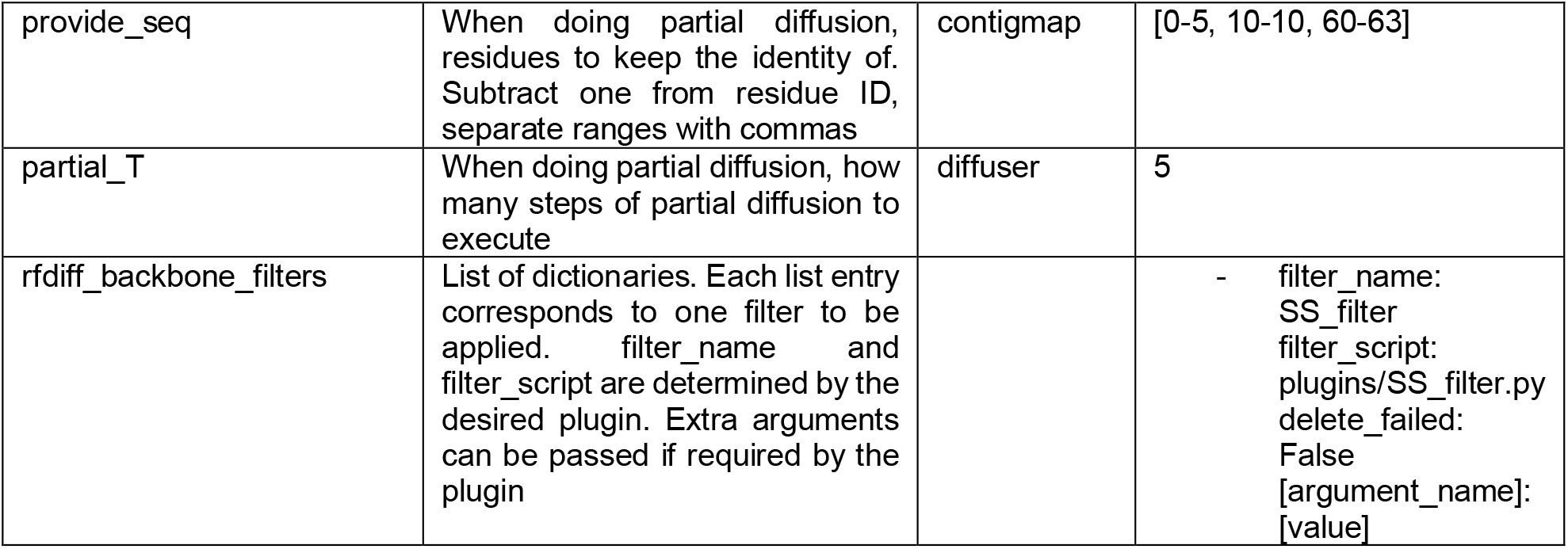
Important parameters for Prosculpt. The list of possible parameters to be passed to RFdiffusion and ProteinMPNN is not exhaustive. Some parameters need to be in dictionary variables which then get passed to individual programs; when this is the case, “dictionary name” is the name of that dictionary. For clarification, look at the included examples.

## Notes

https://github.com/ajasja/prosculpt

### Competing Interest Statement

The authors have declared no competing interest.

